# A Transient Metabolic State In Melanoma Persister Cells Mediated By Chemotherapeutic Treatments

**DOI:** 10.1101/2021.02.21.432154

**Authors:** Prashant Karki, Vahideh Angardi, Juan C. Mier, Mehmet A. Orman

## Abstract

Persister cells are defined as the small fraction of quiescent cells in a bulk cancer cell population that can tolerate unusually high levels of drugs. Persistence is a transient state that poses an important health concern in cancer therapy. The mechanisms associated with persister phenotypes are highly diverse and complex, and many aspects of persister cell physiology remain to be explored. We applied a melanoma cell line and panel of chemotherapeutic agents to show that melanoma persister cells are not necessarily preexisting dormant cells or stem cells; in fact, they may be induced by cancer chemotherapeutics. Our metabolomics analysis and phenotype microarray assays further demonstrated that the levels of Krebs cycle molecules are significantly lower in the melanoma persister subpopulation than in the untreated bulk cell population due to increased utilization rates in persisters. Our data indicate that this observed metabolic remodeling is transient, as the consumption rates of Krebs cycle metabolites are significantly reduced in the progenies of persisters. Given that the mitochondrial electron transport chain (ETC) is more active in the persister subpopulation than in the bulk cancer cell population, we also verified that targeting ETC activity can reduce melanoma persistence. The reported metabolic remodeling feature seems to be a conserved characteristic of melanoma persistence, as it has been observed in various melanoma persister subpopulations derived from a diverse range of chemotherapeutics. Elucidating a global metabolic mechanism that contributes to persister survival and reversible switching will ultimately foster the development of novel cancer therapeutic strategies.

## INTRODUCTION

Conventional cancer therapies target the mechanisms underlying the rapid growth of tumor cells. However, these therapies are usually inefficient for small subpopulations of persister cancer cells that are in a transient “persistence state”.^1–3^ This phenomenon resembles bacterial persistence, which is characterized by slow growth coupled with the ability to tolerate unusually high levels of drugs and has been documented across multiple tumor cell lines and in response to a variety of therapeutic challenges.^1–3^ The molecular mechanisms underlying the observed tolerance of persister cells are highly complex and diverse and could be associated with cell dormancy; drug targets inactivation or alteration; increased drug efflux and DNA damage repair activities; cell death pathway inhibition; and cancer stemness.^4–11^ Persisters are an important health concern. While persistence is defined as a transient, nonmutagenic state, it can serve as a source of drug-tolerant mutants.^3^ Persisters are also thought to underlie the proclivity of recurrent cancers to relapse.^2,6,12^ Recurrence is seen in many tumor types, including skin, lung, pancreas, bladder, and breast cancers, and continues to be a major challenge in cancer therapy.^13^ For instance, a study performed over a period of 20 years (1994–2014) at the Beth Israel Deaconess Medical Center showed that patients with melanoma have an estimated 41.1% recurrence rate.^14^

Unfortunately, melanoma is the most fatal form of skin cancer, and its incidence rate in the U.S. has tripled over the past decade.^15^ The American Cancer Society estimated approximately 106,110 new cases and 7,180 deaths related to melanoma in 2021.^16^

Metabolic reprogramming, including rapid ATP generation, increased biosynthesis of macromolecules, and maintenance of cellular redox balance under nutrient-depleted conditions and other stresses, is one of the hallmarks of cancer^17^ and occurs to meet the essential needs of cancer cells. Aerobic glycolysis, known as the Warburg effect, is the most common feature of metabolic reprogramming observed in cancer cells. This phenomenon is characterized by the increased consumption of glucose via glycolysis and the downregulation of oxidative phosphorylation irrespective of oxygen availability and mitochondrial activity.^18–20^ This shift seems to be essential for supporting the large-scale biosynthetic processes that are required for active cell proliferation.^19,21^ Although aerobic glycolysis appears to occur in many rapidly dividing mammalian cells, this may not necessarily be the case in persisters, which exist in a slowly proliferating state.^22–26^ Metabolic rewiring in persister cells potentially extends beyond glycolysis, and these cells can rely on different metabolic pathways to evade drug effects. Understanding the metabolic state of persisters will provide important insights that are likely to aid the development of novel and broadly effective cancer treatments. A recent study by Hangauer *et al.*^2^ presented an example of the therapeutic promise of targeting persister metabolism. Specifically, the study revealed the existence of a common survival mechanism mediated by the lipid hydroperoxidase GPX4 in persister cell populations derived from breast, melanoma, lung, and ovarian cancers. The team screened a diverse collection of compounds and found that two GPX4 inhibitors (RSL3 and ML210) were selectively lethal to persisters. In a separate study, Shen *et al.* ^27^ revealed the existence of a metabolic mechanism, characterized by the upregulation of fatty acid oxidation, in the melanoma persister cell population mediated by BRAF and MEK inhibitors. Although many studies have shown that oxidative stress plays a critical role in persistence,^11,22,26,27^ we first need to obtain a comprehensive understanding of the metabolic state of persister cells to explore their metabolism as a therapeutic target. We still need to elucidate (i) whether the metabolic rewiring observed in persister cells is a hallmark of cancer persistence, (ii) whether it is a transient state induced by cancer therapeutics and (iii) whether it depends on drug type, concentration and treatment duration.

Our goal in this study was to evaluate the role of metabolism in melanoma persistence and the utility of targeting persister metabolism as a therapeutic strategy. Conventional chemotherapy is one of the most common treatment strategies used to rapidly kill proliferating cancer cells. Unlike targeted therapeutics, chemotherapeutics may not be cancer type specific. However, according to American Cancer Society, chemotherapy is not often used for melanoma patients, as the cancer, upon the conclusion of the treatment, usually starts growing again within several months.^28^ Chemotherapeutics may stimulate a persistence state in melanoma cells, which remains to be characterized. Most chemotherapeutics cause DNA damage, which induces the phosphorylation of ATM and ATR kinases.^29–32^ ATM-mediated growth arrest can be facilitated by the transcription factor p53, which activates the cyclin-dependent kinase (CDK) inhibitor p21.^33,34^ In the absence of functional p53, ATM and ATR can still induce cell cycle arrest, as these regulators, together with CHK1 and CHK2, reduce CDK activity, thus resulting in cell dormancy via the inactivation of cell proliferation-related signaling pathways.^32^ As we think persister cells are potentially induced by chemotherapeutics, the metabolic rewiring associated with growth arrest is inevitable during drug treatment. To further demonstrate whether the induction of growth arrest by chemicals is observed in skin cancer, we analyzed the Library of Integrated Network-Based Cellular Signatures (LINCS) Consortium database.^35,36^ This database, which includes more than 100,000 gene expression profiles, was generated through a data processing pipeline that captures raw data for more than 950 transcripts and infers the expression of nonmeasured transcripts for each cell line and chemical treatment condition.^35^ Our bioinformatics analysis indicates that chemical treatments decrease the expression of cyclins, CDKs, and other important proteins that mediate cell division (e.g., CDK1, CCNB1, CCNB2, CCNA2, CDC25B, CDC20) in skin cancer (Supp. Table 1), which is in agreement with our argument.

In this study, our characterization of the metabolic mechanisms of melanoma persister cells revealed that (i) metabolic rewiring associated with increased mitochondrial activity is a general characteristic of melanoma persisters, (ii) the observed metabolic state in persisters is transient, and (iii) this metabolic state is a result of the inhibition of cell growth, which can be mediated by a wide range of chemotherapeutics.

## RESULTS

### Persistence is represented by a slow-growing cell state, largely induced by chemotherapeutics

In this study, persister subpopulations were derived from gemcitabine (GEM)-treated A375 melanoma cell cultures (Figure 1a). The A375 cell line has BRAF V600E mutations, leading to excessive cellular proliferation and differentiation and increased cell survival.^11,27,37^ BRAF is a protooncogene encoding a serine/threonine kinase of the RAF family.^38^ GEM is a nucleoside that is an analog of deoxycytidine.^39^ Similar to major chemotherapeutic agents, GEM inhibits DNA replication by incorporating itself at the end of the elongating DNA strand. As persister cells have the ability to tolerate high concentrations of drugs, we treated A375 cells with GEM for 3 days to generate a concentration vs. survival ratio profile (Figure 1b); the results showed that the cell survival ratio did not change significantly at concentrations higher than 10 x the half minimal inhibitory concentration (10xIC_50_= 20 nM, see Supp. Table 2).^40,41^ After 3 days of GEM treatment, we gently detached the cells from the flasks, resuspended them in fresh, drug-free growth medium, and incubated them for 24 h to remove dead/late apoptotic cells and collect the persister cells. As shown in the microscope images from the live/dead (blue/green) staining assay (Figure 1c), nearly all persister cells were viable. In this assay, the blue probe stained the nuclei of all cells, and the green probe only stained the nuclei of cells with compromised membranes (see Materials and Methods). Furthermore, an annexin-V fluorescein isothiocyanate (FITC)/propidium iodide (PI) assay^42^ was performed to detect apoptotic cells. One of the early markers of apoptosis is the appearance of phosphatidylserine (PS) on the surface of the cells. PS is usually located in the membrane leaflets that face the cytosol or the cytoplasmic face. However, during apoptosis, PS is exposed on the outer leaflet of the cell membrane.^43^ Annexin V binds to PS with high specificity in the presence of calcium.^42^ Based on staining in this assay, cells were gated into four categories: (i) FITC-negative and PI-negative, (ii) FITC-positive and PI-negative, (iii) FITC-positive and PI-positive and (iv) FITC-negative and PI-positive, representing healthy, early apoptotic, late apoptotic and dead cells, respectively. The data showed that the apoptosis levels in both the parental and surviving persister cell populations were nonsignificant (Figure 1d). Unlike drug-resistant mutants, the progenies of persisters are susceptible to cancer drugs; this phenomenon has been demonstrated in many other studies.^1,2,27^ Therefore, cells surviving GEM treatment were transferred to fresh medium, resuspended and retreated with GEM to verify the transient state of melanoma persister cells (Figure 1a). To demonstrate that persister cells are largely induced by GEM treatment, we performed a cell proliferation study using carboxyfluorescein succinimidyl ester (CFSE) dye, which can freely diffuse across the cell membrane and produce a stable fluorescent signal following an enzymatic reaction with cellular esterases.^44^ For this assay, the prestained cells were treated with GEM or left untreated (control), and the cell proliferation rates of these groups were measured by monitoring the fluorescence dilution rate over time with a flow cytometer. Our results revealed ongoing cell division in the control groups, as evidenced by a reduction in the fluorescent signals, whereas the fluorescent signal was maintained in the treatment groups largely due to a lack of cell proliferation (Figure 1e). The mean fluorescence intensity for the first 3 days for each group was integrated into the fluorescence decay equation to calculate the half-life of the fluorescent signal, further showing that cells treated with GEM grew significantly slower than untreated control cells (Supp. Figure 1). Our microscope images further showed that the surviving cell populations seemed to be heterogeneous, similar to the untreated control cells. We detected preexisting, stem-like spherical cells with high levels of green fluorescence that survived the drug treatment due to the lack of cell proliferation (Figure 1f). However, some surviving cells arose from cell subpopulations with low levels of green fluorescence (Figure 1f), suggesting that they may have been proliferating before drug treatment (Supp. Figure 2); this finding indicates that dormancy can be induced by anticancer drugs (Supp. Figure 2). Since dormancy is often related to cancer stem cells (CSCs), we performed an ALDEFLUOR assay^45,46^ to verify whether persisters are preexisting CSCs. High aldehyde dehydrogenase (ALDH) activity has been verified to be a functional biomarker of melanoma stem cells.^47–49^ We measured the ALDH activity of GEM-treated persister cells as well as the untreated control cells and found no significant increase in ALDH activity in persister cells compared to control cells (Figure 1g), indicating that GEM persisters may represent a subpopulation of cells with a distinct phenotype.

**Figure 1:**
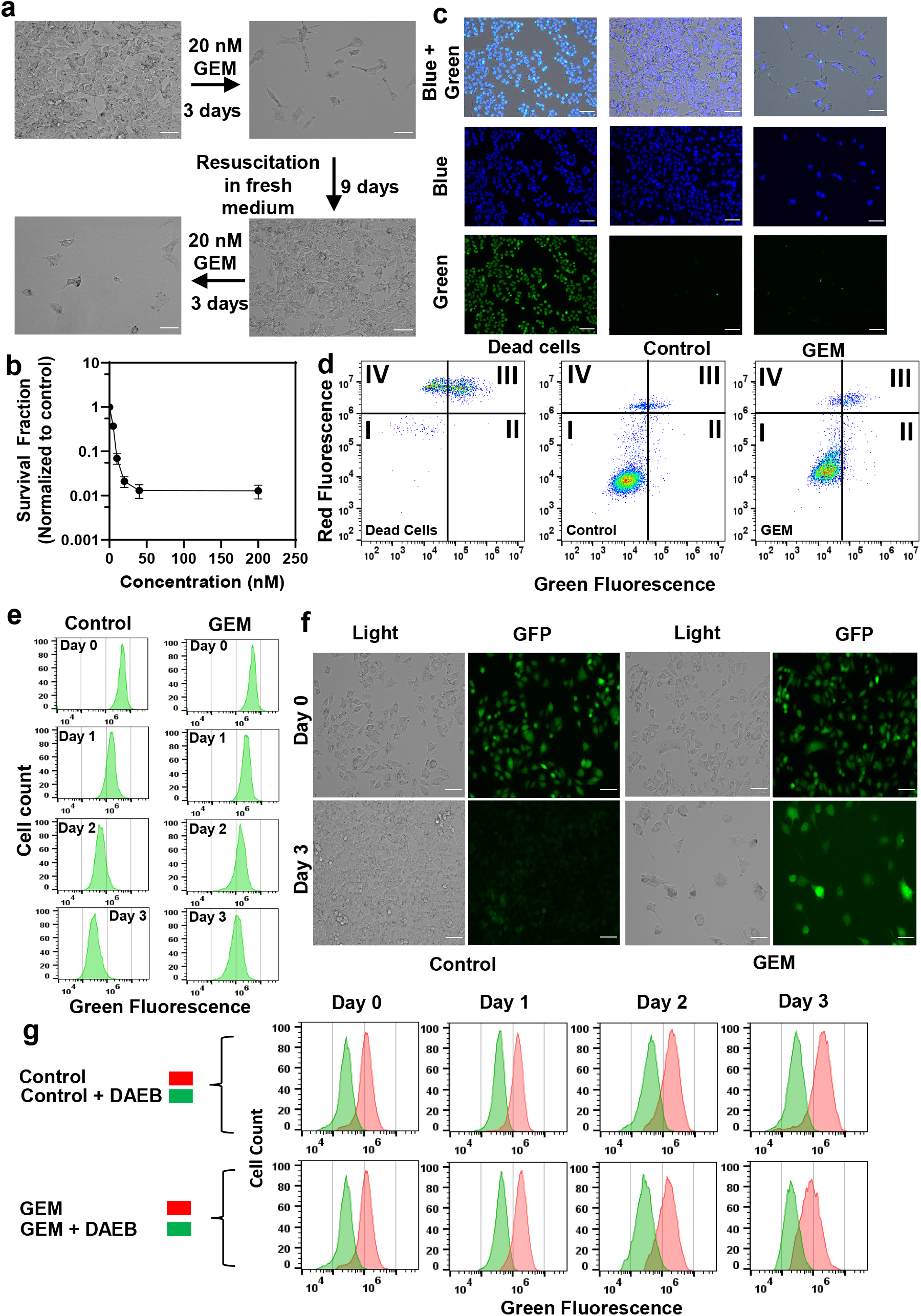
Isolating drug-tolerant persister cells. a) A375 melanoma cells were treated with GEM (10xIC_50_=20 nM, see Supp. Table 2) for 3 days. After the treatment, cells were allowed to recover in fresh, drug-free growth medium and then treated with GEM after recovery to demonstrate the sensitivity of the daughter cells to GEM. Scale bar: 100 μm. b) A375 cells were treated with GEM at the indicated concentrations for 3 days. The surviving cells after the treatment were collected and transferred to fresh medium without GEM. The following day, cell viability was assessed by trypan blue staining (see Materials and Methods). Survival fractions were calculated by normalizing the surviving cell numbers to those in the untreated control groups. The number of biological replicates (N)=4. c) The cells surviving after GEM treatment were collected and transferred to fresh medium without GEM. The following day, cells were stained with ReadyProbes Cell Viability Imaging dyes to assess live (blue) and dead (green) cells. Dead cells were generated by treating the cells with 70% ethanol for 30 min. “Control” represents the live cells that did not receive GEM treatment. Scale bar: 100 μm. d) Cells after GEM treatment were collected and transferred to fresh medium without GEM. The following day, cells were stained with FITC-annexin-V conjugate and PI to detect apoptotic cells. The quadrants of this graph represent (I) live (FITC^−^/PI^−^), (II) early apoptotic (FITC^+^/PI^−^), (III) late apoptotic (FITC^+^/PI^+^) and (IV) dead (FITC^−^/PI^+^) cells. e-f) Melanoma cells prestained with CFSE dye were treated with GEM or left untreated (control), and their fluorescence intensity was monitored at the indicated time points with flow cytometry (E) or florescence microscopy (F). N=4. Scale bar: 100 μm. g) Melanoma cells were treated with GEM or left untreated for 3 days. Every day, the ALDH activity of the cells was assessed with an ALDEFLUOR assay and a flow cytometer. Cells treated with the ALDH inhibitor 4-(dimethylamino)benzaldehyde (DAEB) served as negative controls.

### Persister cells have an altered metabolic state

Tumor cells undergo metabolic alteration to fulfill the energy requirement, sustain the high rate of cell proliferation, avoid the action of therapeutics, improve the overall survivability of the tumor cells, and facilitate other processes.^50^ As persistence is a transient state, we expected that persister cells would undergo metabolic alterations due to their slow or nonproliferating cell state. To identify such metabolic mechanisms, we conducted untargeted metabolomics analysis of GEM-treated cells and untreated control cells. Persisters were generated by treating the cells with 10xIC_50_ of GEM, and untreated cells were generated by culturing the cells in drug-free growth media. In our study, we measured 689 different metabolites that are part of the superpathways involving the following factors: amino acids, peptides, carbohydrates, energy, lipids, nucleotides, cofactors/vitamin and xenobiotics (Supp. Table 3). Each superpathway was further subdivided into 102 subpathways. The obtained MS data were normalized based on protein concentration and statistically analyzed with ANOVA (P≤0.05). For the 102 subpathways, pathway enrichment analysis showed that 57 of them had metabolites that were significantly altered in the persister subpopulation versus the control cells (Supp. Table 3). Unsupervised hierarchical clustering of the metabolic data of four independent biological replicates of untreated or GEM-treated samples reveals a distinct metabolic rewiring taking place in persister cells (Figure 2a). While metabolites associated with dipeptides, phospholipids, sphingosines, the urea cycle, gamma-glutamyl amino acid, ceramides, polyamines, tryptophan, sterols, endocannabinoid, phosphatidylcholines (PC), lysophospholipids and sphingomyelins were upregulated, those associated with glycine, serine, threonine, pentose sugars, vitamin B6, glutamate, the Krebs cycle and branched-chain amino acids (BCAAs) were significantly downregulated (Figure 2a-f, Supp. Table 3).

**Figure 2:**
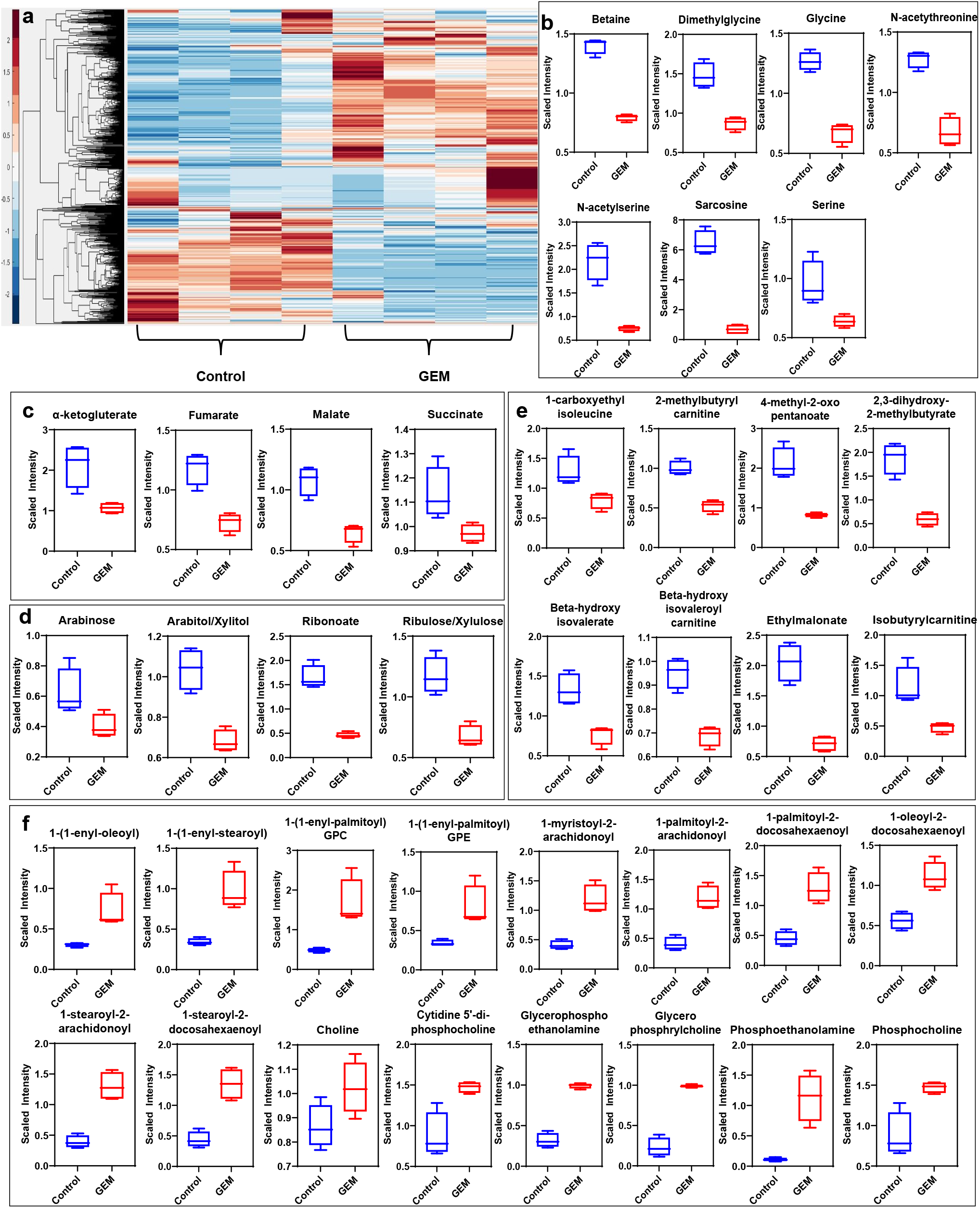
Metabolic alterations in GEM persisters. a) GEM-treated cells and untreated cells were collected for MS analysis to measure their metabolite contents. Unsupervised clustering of the metabolomics data was performed with the Clustergram function in MATLAB. The generated heat maps show metabolite clusters that are upregulated (red) or downregulated (blue) in the treated group compared to the untreated control group. Each column represents a biological replicate; each row represents a metabolite. N=4. b-f) Box plots show the metabolites from the one-carbon metabolism pathway (b), the Krebs cycle (c), the PPP (d), the BCAA metabolism pathways (e), and the lipid metabolism pathway (f) that are significantly altered in GEM persisters compared to control cells. Statistical significance was assessed with ANOVA (P≤0.05). N=4.

Alteration of the lipid metabolism of cancer cells compared to that of nonmalignant cells is a well-studied phenomenon.^51^ This metabolic reprogramming has been shown to be highly dependent on the cancer type and stage. Glycerophospholipids (e.g., PC, PS, phosphatidylethanolamine, (PE), phosphatidylglycerol (PG), phosphatidylinositol and phosphatidylinositol-phosphates (PIPs)) are predominant components of the cell membrane that can play an important role in persistence by modulating the expression and activity of multidrug resistance pumps.^52^ Our metabolomic analysis indicated that phospholipids, particularly PCs, were upregulated in persister cells (Figure 2f). Sphingolipids are another family of membrane lipids known to play a role in the regulation of cell proliferation, apoptosis, migration and inflammation.^53^ Our results show that these metabolites and their associated structural elements (ceramide and sphingosine) are considerably upregulated in persister cells compared with untreated cells (Figure 2f). One-carbon metabolism, as an indicator of the cell nutrient status, functions in the biosynthesis of nucleotides as well as the maintenance of the redox and methylation states required to support the high rate of proliferation in cancer cells.^54^ Our metabolomics results show that metabolites involved in one-carbon metabolism (e.g., glycine, serine, and methionine) were distinctively downregulated in GEM-treated cells (Figure 2b).

Cancer cells overexpress amino acid-degrading enzymes to increase their energy production and to provide metabolites for their anabolic processes. BCAAs, including leucine, isoleucine and valines, are a class of amino acids and were significantly downregulated in GEM-treated cells (Figure 2e). BCAAs are expected to be upregulated in normal cancer cells, as they can be used for various processes such as protein synthesis and energy production.^55^ Of the carbohydrate family, only the pentose phosphate pathway (PPP) was significantly downregulated in GEM-treated cells (Figure 2d). Similar to one-carbon metabolism, the PPP was shown to be important for tumor cells in terms of nicotinamide adenine dinucleotide phosphate (NADPH) production, which is essential for fatty acid synthesis and reactive oxygen species detoxification.^56^ The PPP is tightly interconnected with glycolysis and the Krebs cycle, as they share a number of intermediates, including glucose-6-phosphate (G-6-P), pyruvate and acetyl-CoA. The Krebs cycle is also closely linked to BCAA metabolism, as alpha-ketoglutarate is essential for BCAA metabolism. Our untargeted metabolomics analysis showed that, similar to BCAA and PPP metabolism, the Krebs cycle was downregulated in melanoma persisters, as its intermediates (e.g., alpha-ketoglutarate, fumarate, malate, and oxaloacetate) were significantly less abundant in melanoma persisters than in the untreated bulk cell population; however, we did not observe a significant change in glycolysis intermediates, except for pyruvate, in either group (Figure 2c). Although our MS analysis shows a metabolic alteration in energy metabolism in persister cells, MS does not directly measure intracellular reaction rates (e.g., mitochondrial activity); such measurements are necessary to verify whether metabolic rewiring occurs in persisters.

### Persister cells have increased mitochondrial activity

Cancer cells are known to use aerobic glycolysis to generate substrates for the anabolic processes needed to support cell proliferation.^18,19^ We think that, due to their nonproliferating cell state^1,2^, persisters utilize oxidative phosphorylation rather than aerobic glycolysis. This metabolic rewiring may explain the low levels of Krebs cycle intermediates observed in persisters, as the depletion of these substrates is potentially due to faster consumption of these compounds in persisters. To verify this, we measured the mitochondrial activity of the persister cells using MitoPlates (Biolog Inc., Hayward, CA). Mitoplate assays employ a modified version of tetrazolium dye that can be reduced intracellularly by ETC activity across the membranes of metabolically active mitochondria, resulting in the production of water-soluble formazan. The color change associated with formazan production can be detected by absorbance measurements (OD_590_) and correlates with cellular ETC activities. For this assay, 30 different substrates associated with glycolysis and the Krebs cycle were screened in a 96-well format. A kinetic graph was generated to illustrate the consumption of each substrate by measuring the color intensity of the tetrazolium dye present in each well. The obtained data were then clustered (unsupervised) to generate a heat map (Figure 3a) for all the substrates being studied. Of all the substrates that were tested, persister cells had a higher rate of consumption of Krebs cycle metabolites (specifically, the consumption rates of malate, fumarate and succinate) than untreated cells. Mitoplate screening is a high-throughput assay with limited control over the concentrations of substrates in microarrays. As the concentrations of the substrates were not disclosed, to verify the observed results, these assays were repeated in a generic 96-well plate where the metabolites (i.e., malate, fumarate and succinate) were added manually to achieve a final concentration of 4 mM. The data generated from these modified assays were in agreement with our MitoPlate data (Figure 3b-d).

**Figure 3:**
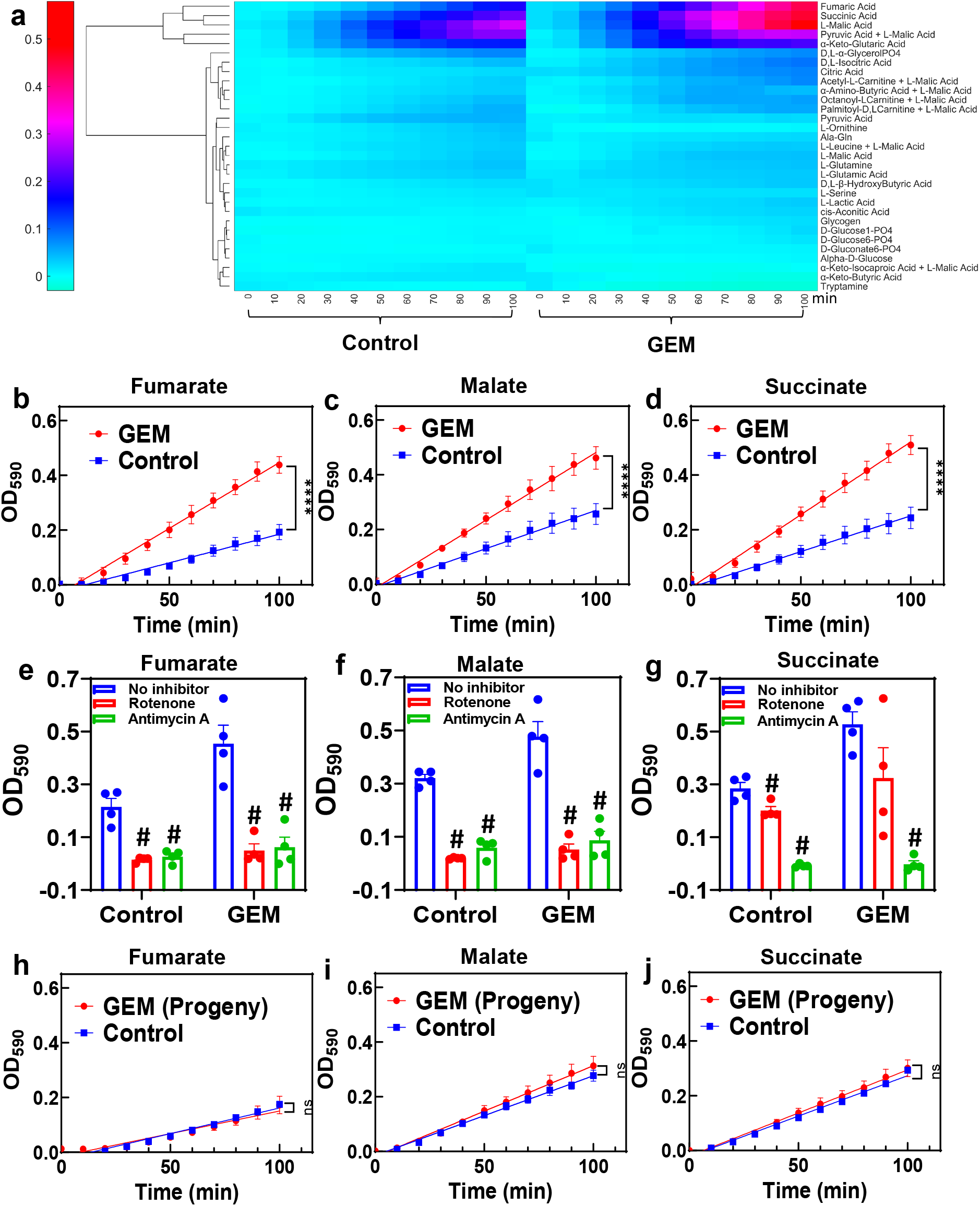
Transient upregulation of Krebs cycle activity in GEM persisters. a) Phenotype microarrays were used to assess the mitochondrial activities of GEM persister cells; 3 × 10^4^ persister or untreated (control) cells were transferred to each well of a phenotype microarray that also included a substrate and tetrazolium-based dye. The consumption rates of substrates were monitored by measuring the OD_590_ at the indicated time points (for 100 min total). Unsupervised clustering of absorbance data was performed using MATLAB. N=4. b-d) The consumption rates for the selected substrates (4 mM) (fumarate, malate and succinate) were measured with the modified MitoPlate assay (see Materials and Methods). Linear regression analysis was performed using F statistics using GraphPad Prism (**** P<0.0001). N=4. e-f) Bar graphs represent the OD_590_ data of the modified MitoPlate assays for fumarate, malate and succinate and indicate their consumption rates at 100 min in the presence of rotenone (10 μM) or antimycin A (10 μM). N=4. Statistical significance was assessed by performing pair-wise t-test. # represents a significant difference between the inhibitor (rotenone or antimycin A) and “no inhibitor” groups (P<0.05). h-j) Persister cells were transferred to fresh medium without GEM to stimulate resuscitation. After the third passage, the daughter cells were collected, and their consumption rates for fumarate, malate and succinate were measured with the modified MitoPlate assay. Untreated parental cells were used as a control. N=4; ns: the slopes are not significantly different.

To further verify the accuracy of the assays, control experiments, in which ETC complexes were inhibited with rotenone and antimycin A, were conducted. Rotenone is a complex I inhibitor, and antimycin A inhibits complex III of the ETC.^57,58^ Therefore, the substrates capable of producing only NADH (i.e., malate and fumarate) and the substrates producing both NADH and FADH_2_ (i.e., malate, fumarate and succinate) should not give any absorbance reading in the presence of rotenone and antimycin A, respectively, in modified MitoPlate assays. The data generated support our argument and validate the efficacy of the MitoPlate assay (Figure e-g). Finally, as persistence is a temporary state, the observed metabolic alteration should also be transient. Cells that survived GEM treatment were collected and regrown in fresh growth medium. After 9 days of resuscitation, the cells resumed their growth cycle and started to proliferate (Figure 1a). The progenies of the resuscitated cells after the 3^rd^ passage were collected to assess their mitochondrial activity. As expected, the consumption rates of malate, fumarate and succinate in persister progenies were similar to those of untreated control groups (Figure 3h-j).

### The observed metabolic rewiring is independent of GEM concentration and treatment time

The treatment duration and chemotherapeutic concentration can play a significant role in persister cell metabolism. To assess the effect of the treatment period, A375 cells were treated with GEM (10xIC_50_) for 9 days, and the surviving cells were collected for MitoPlate assays, which showed that the consumption rates of Krebs cycle substrates were still higher in GEM-treated cells than in untreated control cells (Supp. Figure 3a). However, interestingly, the control cells cultured for 9 days had higher consumption rates of Krebs cycle substrates than the control cells cultured for 3 days (Figure 3a and Supp. Figure 3a). This observation might be due to the senescence associated with the late stationary-phase cells cultured 9 days, which is consistent with our central argument. To assess the effects of drug concentrations on persister metabolism, we isolated persisters by treating the cells with 100xIC_50_ GEM for 3 days (Supp. Figure 3c-d). Similar to the 10xIC_50_ treatment results, the surviving cells exhibited higher rates of consumption for Krebs cycle substrates than untreated cells (Supp. Figure 3b and 3e-g). Persister cells obtained by 100xIC_50_ GEM treatment were resuspended for a second round of cell survival and MitoPlate assays, showing that the progenies of persister cells were sensitive to GEM (Supp. Figure 3d), and the observed metabolic alteration was reversible (Supp. Figure 3d). Persister cells obtained from 100xIC_50_ GEM treatment were also viable, grew slowly, and exhibited phenotypic heterogeneity (Supp. Figure 3c), in agreement with the data generated from 10xIC_50_ treatments (Figure 1). Interestingly, the cells surviving 100xIC_50_ GEM treatment required ~32 days to resuscitate. This was significantly longer than the resuscitation period of the 10xIC_50_ treatment group (~9 days), indicating that the resuscitation period is concentration dependent, although increasing the GEM concentration does not affect metabolic rewiring.

### Chemotherapeutic-induced metabolic rewiring is conserved in melanoma persisters

As GEM is a very common chemotherapeutic agent, we wanted to test whether the observed results were also valid for the other chemotherapeutic agents listed in Supp. Table 2. Cytarabine (CYT) is an antimetabolite similar to GEM; camptothecin (CAM), doxorubicin (DOX) and etoposide (ETO) inhibit topoisomerase; cisplatin (CIS) and temozolomide (TEM) is an alkylating agent; vinorelbine (VIN) and paclitaxel (PAC) impair the formation of spindle fibers; and mitomycin-c (MIT) induces cross-linking of DNA.^59^ Cells were treated with these therapeutic agents at 10xIC_50_ doses (Supp. Table 2), except TEM, which was applied at 5xIC_50_, as the 10xIC_50_ dose required a high dimethyl sulfoxide (DMSO) solvent content (>1%). Live/dead staining was performed for all treatments to ensure high persister cell viability (Supp. Figure 4). The mitochondrial activity for each treatment was assessed using modified MitoPlate assays, which demonstrated that the chemotherapeutic agents generally increased the consumption rates of Krebs cycle substrates in melanoma cells (Figure 4a). Furthermore, we performed ALDEFLUOR assays for all treatment groups (Supp. Figure 5) to compare their ALDH activities with those of untreated control groups. Similar to the results obtained from the GEM treatment, most treatments did not significantly alter cellular ALDH activity. However, TEM- or MIT-treated cells showed significantly lower ALDH activity than control cells, further supporting that chemotherapeutic persistence may not be directly linked to CSC phenotypes. Our flow cytometry- and microscopy-based cell proliferation assays further showed that cells from all treatment groups had undergone a state of negligible growth, indicating that chemotherapeutic-induced growth arrest is conserved in melanoma cells (Figure 4b, Supp. Figure 1 and Supp. Figure 6). Our microscopy images showed that when compared to the untreated control cells, treated cells had altered morphology with high fluorescence intensity due to the induction of growth arrest (Figure 4b vs. control group in Figure 1f). Overall, the upregulation of Krebs cycle activity is conserved in melanoma persister populations derived from various chemotherapeutic treatments, despite the diverse morphological changes observed in these persister populations (Figure 4b and Supp. Figure 4).

**Figure 4:**
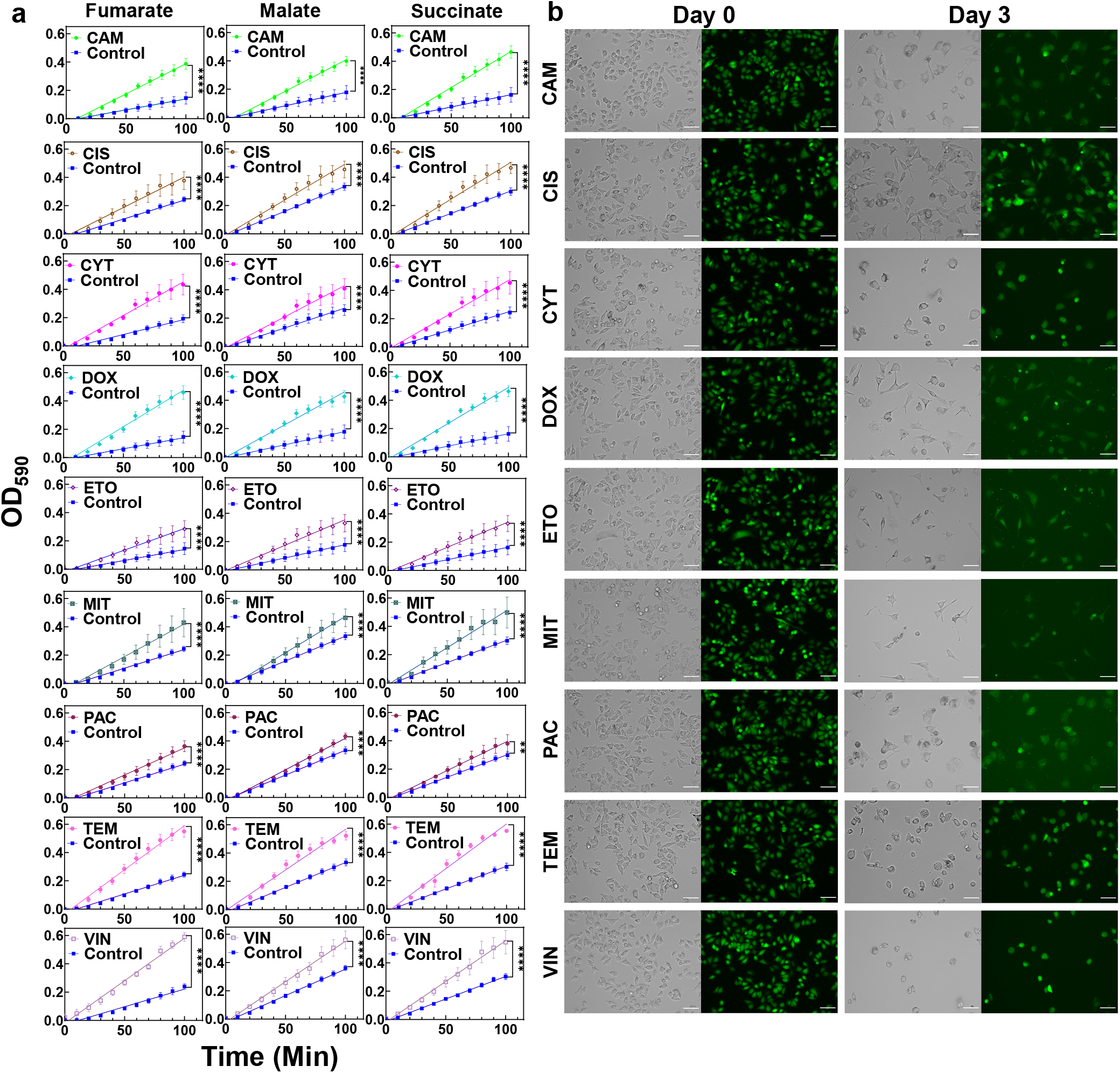
Chemotherapeutic treatments induced slow growth and enhanced Krebs cycle activity. a) Melanoma cells were treated with the indicated chemotherapeutic agents for 3 days. After treatments, the cells were transferred to fresh growth medium and incubated for 24 h. Then, the cells were collected for modified MitoPlate assays to measure the consumption rates of Krebs cycle substrates. The concentrations of chemotherapeutic agents were 10xIC_50_, except TEM, whose concentration was 5xIC_50_ (Supp. Table 2). Linear regression analysis was performed using F statistics using GraphPad Prism (**** P<0.0001). N=4. b) Melanoma cells stained with CFSE were treated with the indicated agents for 3 days as described above. Prior to treatment (day 0) and after treatment (day 3), the green fluorescence of the cells was monitored by florescence microscopy. Scale bar: 100 μm.

### Cotreatment with ETC inhibitors can compromise persister survival

We used a microarray plate containing mitochondrial inhibitors (meclizine, rotenone, pyridaben, berberine, malonate, carboxin, alexidine, antimycin A, myxothiazol, phenformin, trifluoromethoxy carbonylcyanide phenylhydrazone (FCCP), 2,4-dinitrophenol, diclofenac, valinomycin, calcium chloride, celastrol, gossypol, nordihydroguaiaretic acid, trifluoperazine (TFZ), polymyxin B, amitriptyline and papaverine) to test the effects of these inhibitors on cell viability with or without GEM. The chemical library had four concentrations of each inhibitor, but these concentrations were not disclosed by the company (Biolog, Inc.). As we wanted to identify a chemical compound that is selectively and effectively lethal to GEM persisters, we focused on the wells with the lowest inhibitor concentrations (Figure 5a), and our observations revealed that TFZ might be a potential chemotherapeutic adjuvant (Figure 5a). Although a number of inhibitors, including glossypol, valinomycin, and celastrol, were found to be effective at higher concentrations, these inhibitors were also lethal to cancer cells in the absence of GEM (Supp. Figure 7). TFZ falls under the class of antipsychotic drugs known as phenothiazines, which have been shown to enhance the cytotoxic effects of chemotherapeutic agents.^60^ These drugs have also been shown to inhibit tumor progression by interacting with various cell signaling pathways, such as Wnt, MAPK, Akt, and ERK and by inhibiting drug efflux pumps.^61^ To determine whether inhibition of persisters is a more general characteristic of phenothiazines, we tested two additional FDA-approved phenothiazine drugs, thioridazine (TDZ) and fluphenazine (FPZ), that were not in our drug screen. Notably, TDZ was recently demonstrated to impair melanoma tumor progression in an animal model.^27^ Although all three phenothiazine inhibitors reduced the cell survival fractions across a wide range of concentrations when tested with GEM (Figure 5b-d), these inhibitors (at concentrations greater than 10 μM) could also kill the cancer cells in the absence of GEM (Figure 5b-d). TFZ was also found to be effective in the presence of most of the chemotherapeutics at the indicated concentrations (Figure 5e). TEM, which is used most often for melanoma patients, has become very effective in the presence of TFZ, as the level of survived cells in cotreatment cultures is around the limit of detection (Figure 5e). Although we did not test a wide range of drug concentrations and analyze the synergetic interactions between the drugs in these cotreatments, our results suggest that metabolic inhibitors can potentially boost the effectiveness of the existing chemotherapeutic drugs.

**Figure 5:**
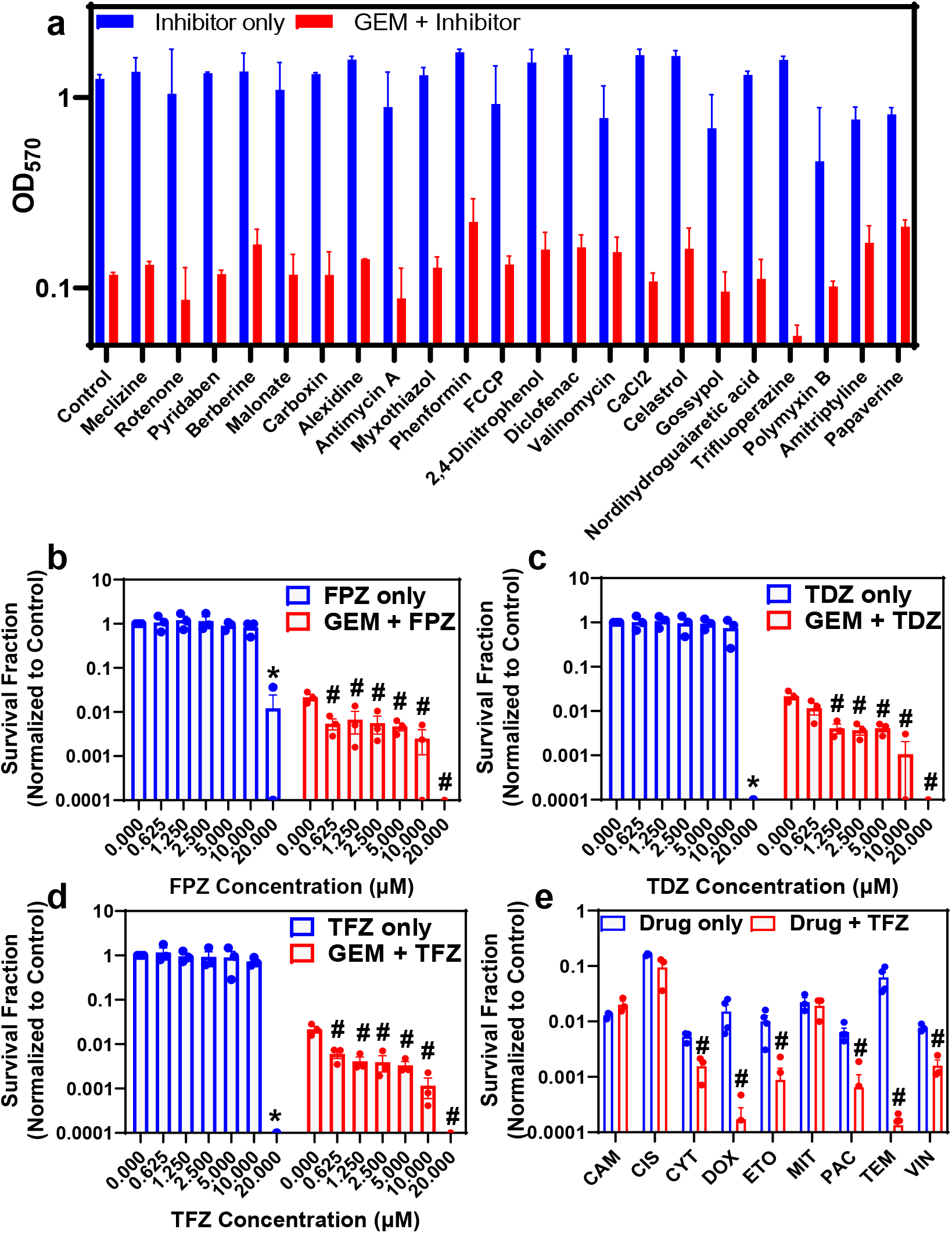
Cotreatment with ETC inhibitors reduces persister survival. a) Melanoma cells incubated in fresh growth medium in a 96-well plate for 24 h were treated with various ETC inhibitors (blue) or cotreated with GEM and ETC inhibitors (red) for 3 days. After the treatment, the media in the wells were replaced with fresh drug-free media. After 24 h of incubation, the MTT assay was conducted to assess cell viability by measuring the absorbance (OD_570_) of all tested combinations with a plate reader. N=2. b-d) Melanoma cells were treated with GEM (10×IC_50_) in the presence of TFZ, TDZ and FPZ at the indicated concentrations for 3 days. After the treatments, the cells were resuspended in fresh drug-free medium and incubated for 24 h. Then, the cell viability was assessed by trypan blue staining using an automated cell counter. * represents a significant difference between the untreated control and inhibitor-only groups, highlighted by blue bars (t-test, P<0.05). # represents a significant difference between the cotreatment and GEM-only groups, highlighted by red columns (t-test, P<0.05). e) Melanoma cells were treated with the indicated chemotherapeutic agents and/or TFZ (10 μM) for 3 days. The concentrations of chemotherapeutic agents were 10xIC50, except TEM, whose concentration was 5xIC50 (Supp. Table 2). After the treatments, the cells were collected and incubated in fresh, drug-free medium for 24 h, and then, the cell viability was assessed with STYO60 (red)/SYTOX (green) dyes using a flow cytometer, as the number of surviving cells in some conditions was under the limit of detection for the automated cell counter. # represents a significant difference between the cotreatment (drug+FPZ) and drug-only groups (t-test, P<0.05).

## DISCUSSION

Conventional chemotherapeutic agents target fast-growing cells such as tumor cells. Because of their slow or no-growth state, persister cell tolerance to treatment has been attributed to cell dormancy.^62,63^ Cancer cell dormancy can be induced by various mechanisms, such as the activation of signaling pathways for autophagy, reactive oxygen species production, and DNA damage repair, that are generally triggered by extracellular stress (e.g., nutrient depletion, hypoxia, overpopulation, therapeutics).^62,63^ Cell dormancy is regulated by many external and internal factors via a highly integrated signaling network and is one of the most common phenotypic states observed in many drug-tolerant cell types, including CSCs. While CSCs exhibit some persister-like characteristics, not all persisters have CSC biomarkers, as shown in our results, indicating that persisters may represent a subpopulation with a distinct phenotype. While persister cells were shown to have CSC biomarkers in previous studies,^2,6^ it is not clear to us whether these biomarkers were induced by the targeted therapeutics used in those studies. Our results indicate that chemotherapeutics may facilitate a transient persistence state, which may lead to the downregulation of anabolic pathways (due to the observed growth arrest), thus diverting glycolytic intermediates to the Krebs cycle, the most efficient energy-producing pathway. This also explains why the abundance of glycolytic metabolites was not altered in persister cells despite the significant alterations in the abundance of Krebs cycle metabolites.

In this study, we first conducted untargeted metabolomics analysis to identify the metabolic pathways that were significantly altered in GEM-treated persister cells compared to control cells. These pathways included the lipid metabolism, BCAA metabolism, one-carbon metabolism, and the Krebs cycle and the PPP. From the lipid superpathways, PC, PE, sphingosines, ceramides and lysophospholipids were primarily upregulated in GEM-treated cells. The accumulation of PC due to overexpression of lysophosphatidylcholine acyltransferase 2 (LPCAT2) can induce drug tolerance in cancer cells, as reported by Cotte *et al*.^64^ The study revealed that LPCAT2 increases the resistance of cancer cells to immunogenic cell death and mediates chemoresistance by promoting the antiapoptotic response to endoplasmic reticulum stressors.^52,64^ Ceramides have been reported to have dual functions in drug resistance. They can induce either chemosensitivity or chemoresistance depending on the structure and length of their fatty acyl chains.^65^ BCAA metabolism, which involves essential amino acids, such as valine, leucine and isoleucine, has been studied extensively in cancer cells.^66,67^ It is closely linked to the Krebs cycle, as alpha-ketoglutarate is needed to initiate the degradation of valine, isoleucine and leucine.^66^ Studies have shown that enzymes that catalyze the first step of BCAA degradation, branched-chain aminotransferase 1 (BCAT1) and branched-chain aminotransferase 2 (BCAT2), are commonly upregulated in cancer cells. BCAT1 in particular is associated with cancer cell growth and has been proposed as a prognostic cell marker.^66,68,69^ In addition, many studies have explored BCAT1 as a potential target for cancer therapeutics, as it is also linked to cell proliferation via m-Torc1 activity.^70^ BCAA metabolism has been shown to alter gene expression in cancer cells by altering the epigenome. Epigenetic changes can affect several cellular processes that can induce drug tolerance in cancer cells. A recent study by Wang *et al.* showed that H3K9 demethylation-mediated epigenetic upregulation of BCAT1 can promote tyrosine kinase inhibitor (TKI) tolerance in epidermal growth factor receptor (EGFR)-mutant lung cancer cells.^68^

The upstream metabolites of the Krebs cycle are needed for the initiation of PPP metabolism. Along with one-carbon metabolism, the PPP regulates NADPH production in cancer cells.^71^ Additionally, it has been hypothesized that slow-growing/drug tolerant cells, such as CSCs, have an increased rate of PPP metabolism compared to the bulk cancer cell population.^72^ Debeb *et al.* showed that PPP metabolism is upregulated in histone deacetylase inhibitor-induced CSCs and is regulated by an increase in the level of G-6-P dehydrogenase, a rate-limiting enzyme in PPP metabolism.^73^ Given that MS does not directly measure intracellular reaction rates, subsequent assays are necessary to link the abundance of metabolites to their turnover rates. Our metabolomics data show that the majority of Krebs cycle metabolites were significantly downregulated in GEM-treated cells due to their increased consumption rates, and these findings were verified by MitoPlate assays. It is known that cancer cells prefer aerobic glycolysis over oxidative phosphorylation; however, as persisters are slow-growing cells, they might not depend on aerobic glycolysis as extensively. This oxidative phosphorylation addiction of drug-tolerant cells has been reported across multiple tumor cell lines and in response to a variety of therapeutic challenges,^10,23–27^ supporting our hypothesis of a conserved, transient metabolic phenomenon mediated by chemotherapeutic treatments. Given that increased oxidative phosphorylation is a potentially conserved characteristic of melanoma persistence, ETC inhibitors can be used as adjuvants for persister therapeutics, as verified by our screening assay of known metabolic inhibitors.

## MATERIALS AND METHODS

### Cell lines and chemicals

The melanoma cell line (A375) was purchased from American Type Culture Collection (ATCC) (Manassas, VA). Unless otherwise stated, all chemicals and growth media were obtained from Fisher Scientific (Waltham, MA). A375 cells were maintained in Dulbecco’s modified Eagle’s medium (DMEM) supplemented with 10% fetal bovine serum (FBS), 100 units penicillin and 100 μg streptomycin/mL at 37 °C in a 5% CO_2_ incubator. MitoPlates, S-1 (catalog# 14105) containing glycolysis and Krebs cycle substrates, and I-1 (catalog # 14104) containing ETC inhibitors were obtained from Biolog, Inc. (Hayward, CA). Saponin (catalog# 47036), used as a cell permeabilization reagent, was purchased from Sigma Aldrich (St. Louis, MO). Stock solutions for all chemotherapeutic agents were prepared with DMSO as the solvent. Mitochondrial inhibitors (TFZ, TDZ and FPZ) were dissolved in sterile deionized (DI) water. The cells were always cultured in DMEM at 37 °C with 5% carbon dioxide (CO_2_) in a humidified incubator; they were treated with chemotherapeutics when they reached a confluency of ~40-50%.

### Transcriptomics dataset analysis

Preliminary inspection included an analysis of the 100 most upregulated and 100 most downregulated genes in melanoma cell lines in the Broad Institute’s Connectivity Map (CMAP) dataset. The CMAP dataset contains information on the mRNA-level changes (in terms of transcript abundance) of a collection of 12,328 human genes after treatment with 118,050 unique perturbation agents and between cell line pairs; these expression data are collectively referred to as signatures. The data were prepared using the L1000 assay, which is used to measure the “Landmark” 978 genes that can, through computational analysis, derive sufficient information about the transcriptional state of a cell. Using this dataset, we derived the 100 most upregulated and 100 most downregulated genes by filtering the z-score data matrix to only include the A375 melanoma cell line. In the new matrix, each row represented the expression level of a gene (defined by z-score) in the melanoma cell line, and each column represented a chemical agent with which the melanoma cell line was treated. The most upregulated genes were selected by counting the number of treatments with a z-score greater than 2 for each gene (P<0.05), and the top 100 genes with the highest count were reported. Likewise, the 100 most downregulated genes were selected by counting the number of treatments with a z-score less than −2 for each gene (Supp. Table 1).

### Persister assays

Approximately 2.5 × 10^6^ cells were suspended in 15 ml of DMEM, plated in T-75 flasks and incubated for 24 h to obtain the desired confluency (~40-50%). Then, the medium was removed and replaced with fresh growth medium containing a chemotherapeutic agent at 10x or 100xIC_50_, as listed in Supp. Table 2. The control cells were treated with the solvent (i.e., DMSO) only. After 3 days of treatment, the cells were washed with 10 ml of Dulbecco’s phosphate-buffered saline (DPBS) twice and detached from the flasks with 2 ml of trypsin-EDTA (0.25% trypsin and 0.9 mM EDTA) for ~1-2 min. After ~1-2 min, 5 ml of DMEM was added, and the cell suspension was transferred to a 10-ml centrifuge tube. The cell suspension was centrifuged at 800 rpm for 5 min, and the supernatants were removed. The cell pellets were resuspended in fresh drug-free media and plated in a T-75 flask. After 24 h of incubation, dead cells floating in the culture medium were removed, and the adherent, live cells were collected for the subsequent assays described below. Of note, when the cells were treated with drugs for 9 days, the medium was changed every 3 days.

To generate kill curves, 3 × 10^5^ cells were plated in each well of a 6-well plate with 3 ml of DMEM and incubated as described above. Similarly, the cells were treated with chemotherapeutics for 3 days and then collected to count the live cells with trypan blue staining^74^ using a countess II automated cell counter (catalog# A27977, Thermo Fisher Scientific). The ratio of surviving cells to untreated control cells was plotted to generate a kill curve profile.

### Live/dead staining

After chemotherapeutic treatments, cells were collected and transferred to fresh medium in a 12-well plate. After 24 h of incubation, the medium with dead cells was removed and replaced with fresh DMEM. Live/dead staining was performed with the ReadyProbes Cell Viability Imaging Kit (Blue/Green) (catalog# R37609, Thermo Fisher Scientific) as described by the protocol provided by the vendor. Fluorescence quantification of stained cells was carried out in standard blue fluorescence (excitation: 360 nm and emission: 460 nm) and GFP (excitation: 470 nm and emission: 525 nm) channels by EVOS M7000 florescence microscopy (catalog# AMF7000, Thermo Fisher). The NucBlue live cell reagent is cell permeant, and the NucGreen dead cell reagent is cell impermeant. Hence, dead cells emit green and blue fluorescence, while live cells only emit blue fluorescence. Live and dead cells were used as controls; dead cells were generated by treatment with 70% ethanol for 30 min.

### Apoptosis

We performed apoptosis assays using the annexin-V FITC/PI kit manual (catalog# P50-929-7; Thermo Fisher Scientific). Cells treated with chemotherapeutics were resuspended in fresh medium and plated in a T-75 flask at 37 °C for 24 h. After 24 h, the cells were collected and resuspended in PBS to obtain a density of 5 × 10^5^ cells per ml. Two hundred microliters of the cell suspension was transferred to a microcentrifuge tube. The cell suspension was centrifuged at 800 rpm for 5 min. The supernatant was removed, and the pellet was resuspended in 195 μl of binding buffer. Five microliters of annexin V-FITC solution was added, and the cell suspension was incubated for 10 min at room temperature in the dark. Following incubation, the washing step was repeated to remove any excess dye. The cell pellet was resuspended in 190 μl of binding buffer and stained with 10 μl of PI for the detection of dead cells. Finally, the cell suspension was transferred to a 5-ml test tube containing PBS to obtain a final volume of 1 ml cell suspension. The sample was analyzed with a flow cytometer. The cells were excited at 488 nm and 561 nm to assess green (annexin V-FITC) and red (PI) fluorescence, respectively. The green fluorescence was detected with a 520 nm emission filter; the red fluorescence was detected with a 615 nm emission filter. Cells that are FITC-positive but PI-negative are in the early phase of apoptosis; cells that are both FITC-positive and PI-positive are in the late phase of apoptosis, and cell that are both FITC-negative and PI-negative are live cells. Untreated live cells, dead cells and cytarabine-treated cells were used to gate the cell subpopulations in flow cytometry diagrams. Dead cells were generated by treatment with 70% ethanol for 30 min. Cytarabine is known to induce apoptosis;^75^ cells were treated with 50 μM cytarabine for 3 days before staining the cells with the dyes.

### Metabolomics study

After 3 days of GEM (10xIC_50_) treatment, the surviving cells were collected in a 10-ml centrifuge tube, washed with 2 ml PBS by centrifugation (5 min at 800 rpm) and pooled in a microcentrifuge tube to obtain ~100 μl of cell pellet. A dry ice/ethanol bath was used to rapidly cool and freeze the cell pellet. Untreated cells were used as a control. The frozen samples were sent to Metabolon Inc. (Morrisville, NC). The sample extraction, instrument settings, and conditions for the MS platforms followed Metabolon’s protocols (see details in article^76^). Initial data analysis was performed by Metabolon. Briefly, the obtained biochemical data were normalized to the protein concentration (assessed by Bradford assay) of each respective sample. The normalized data were used to form a matrix to perform unsupervised hierarchical clustering with the Clustergram function in MATLAB. Metabolites in persisters were compared with those in control groups using ANOVA with a significance threshold of P ≤ 0.05. A Q-value was used to estimate the false discovery rate, and low Q-values (Q < 0.1) indicated high confidence in the results.

### MitoPlate assay

To assess the mitochondrial function of cells, phenotype microarray plates (S-1, catalog# 14105) were used. The assay employed Biolog Mitochondrial Assay Solution (BMAS, catalog# 72303) together with dye mix MC (tetrazolium-based dye, catalog# 74353) provided by Biolog, Inc. In a 50-ml sterile reservoir, 2x BMAS, MC, 960 μg/ml saponin and sterile water were gently mixed in a 6:4:1:1 ratio. Thirty microliters of the assay mixture was distributed to each well of the 96-well microarray and incubated at 37 °C for 1 h to dissolve the preloaded substrates.

Control or chemotherapeutic-treated cells were collected in a 10-ml centrifuge tube and centrifuged at 800 rpm for 5 min. The supernatant was removed, and the cell pellet was washed with PBS twice to remove any debris. Finally, the cell pellet was resuspended in 1x BMAS to achieve a final cell density of 1 × 10^6^ cells per ml. Thirty microliters of the cell suspension was pipetted into each well of the microarray containing the assay mixture. The final assay mixture was composed of 3 × 10^4^ cells per well. After inoculation, the OD_590_ was measured every 10 min with a Varioskan Lux Microplate Reader (catalog# VLBL00GD0, Thermo Fisher Scientific).

### Modified MitoPlate assay

To verify the accuracy of the MitoPlate assay, the same procedure was repeated in a regular 96-well plate with slight modification. Similar to the MitoPlate assay described above, the assay mixture consisted of BMAS, dye and saponin. However, sterile water was replaced with a solution consisting of 96 mM Krebs cycle substrates (i.e., sodium malate, sodium fumarate or sodium succinate). BMAS, MC, saponin and substrate solution were mixed at a 6:4:1:1 ratio, and 30 μl of the assay mixture was transferred to each well of a standard 96-well plate. Similarly, 30 μl of the cell suspension in 1x BMAS was added to each well of the 96-well plate containing the assay mixture so that each well contained 4 mM substrate and 3 × 10^4^ cells. After inoculation, the OD_590_ was measured every 10 min with a microplate reader. For the control conditions, the ETC inhibitors rotenone or antimycin A were added to the assay mixtures. The final concentration of the inhibitors in the culture was 10 μM.

### Cell growth assay

A375 cells were stained with CFSE dye using CellTrace proliferation kits (catalog# C34570, Thermo Fisher Scientific). The cells were stained with 5 μM CFSE dye following the protocol in the manual. A total of 3 × 10^5^ stained cells were seeded in each well of a 6-well plate and incubated for 24 h. After 24 h, the medium was removed and replaced with fresh DMEM with chemotherapeutic agents at the indicated concentrations. The cells were treated for six days, and the growth medium was changed every three days. Every 24 h, cells were detached from the wells with trypsin, collected and resuspended in 1 ml PBS for analysis with a flow cytometer. The cells were excited at 488 nm, and green fluorescence was detected with a 520-nm emission filter. The fluorescence half-life for all conditions was calculated using the decay equation below:

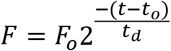

where F_o_ is the mean fluorescence intensity for cells at time t_o_; t_d_ is the half-life time; and F is the mean fluorescence intensity for cells at time t. In this study, t_o_ was chosen as day 0. The half-life time was calculated with SOLVER in Excel by minimizing the sum of normalized mean square errors (NMSE) between experimental and predicted model data.

### Microscopy analysis for cell growth

Cells were stained with 5 μM CFSE dye following the protocol provided in the CellTrace kit. A total of 3 × 10^5^ cells were then seeded in each well of a 6-well plate and incubated for 24 h. After 24 h, the medium was removed from each well and replaced with medium including a chemotherapeutic agent. After 3 days of incubation, cells were washed with 3 ml DPBS, detached with 200 μl of trypsin, resuspended in 1 ml medium, transferred to a 1.5-ml microcentrifuge tube and centrifuged at 800 rpm for 5 min. The supernatant was removed, and the cell pellet was resuspended in 1 ml fresh drug-free growth medium and transferred to each well of a 6-well plate. After 24 h of incubation, the growth medium was replaced with 1 ml DPBS. The cells were then analyzed under an EVOS M7000 fluorescence microscope (excitation: 470 nm and emission: 525 nm).

### ALDEFLUOR assay

An ALDEFLUOR assay kit (catalog# NC9610309, Thermo Fisher Scientific) was used to measure the cellular ALDH activity. A total of 3 × 10^5^ cells were plated in each well of a 6-well plate with 3 ml of DMEM and incubated for 24 h. After 24 h, the growth medium was removed and replaced with fresh growth medium containing 20 nM GEM. Every 24 h, the cells were washed with 3 ml DPBS, detached with 200 μl trypsin, resuspended in 1 ml growth medium and transferred to a microcentrifuge tube. The cell suspension was centrifuged at 800 rpm. for 5 min, and the supernatant was removed. This washing procedure was repeated with DPBS to remove all the residuals. Finally, the cell pellet was resuspended in 1 ml of ALDEFLUOR assay buffer. Five microliters of the activated ALDEFLUOR reagent was added to the cell suspension and mixed. After mixing, 500 μl of the cell suspension was immediately transferred to another microcentrifuge tube containing 5 μl of diethylamino-benzaldehyde (DEAB), which was used as a negative control, as DEAB inhibits ALDH activity. The samples were then incubated at 37 °C for 45 min. After incubation, the cell suspension was centrifuged at 800 rpm. for 5 min, and the supernatant was removed. The cell pellet was resuspended in 500 μl of ice-cold ALDEFLUOR assay buffer and transferred to a 5-ml test tube. Each sample was stained with 1.5 μM PI, incubated for 15 min at room temperature and analyzed by flow cytometry. The cells were excited at 488 nm and 561 nm for green and red fluorescence, respectively. The green fluorescence was detected with a 520-nm emission filter; the red fluorescence was detected with a 615-nm emission filter.

### Inhibitor screening

Approximately 1 × 10^4^ cells were suspended in 150 μl of DMEM, plated in each well of a 96-well plate and incubated for 24 h. Then, the medium was removed and replaced with 150 μl of DMEM containing GEM (20 nM) and/or the ETC inhibitors obtained from the Biolog I-1 plate (catalog # 14104). The ETC inhibitors are in their solid forms in the plate; therefore, 150 μl of DMEM with or without GEM was transferred to each well of the I-1 plate. After 2 h of incubation, the media were transferred from the I-1 plate to the 96-well plate containing the cells, as described above. After 3 days of treatment, the medium was removed, and the cells were washed with 100 μl of DPBS twice. Then, 150 μl of drug and inhibitor-free medium was transferred to each well of the 96-well plate. After incubating the cells for 24 h, the growth medium was removed and replaced with 100 μl of fresh medium. Finally, 10 μl of 3-(4,5-dimethylthiazol-2-yl)-2,5-diphenyltetrazolium bromide (MTT, catalog # 97062-376, VWR) (5 mg/ml) was added to each well to measure the cell viability, and the cells were incubated at 37 °C for 3 h. At 3 h, the medium was removed and replaced with 100 μl of MTT solubilization buffer.^77^ After 20 min of incubation at 37 °C, the OD_570_ was measured with a microplate reader.

### Validation of the inhibitor screening assay results

A total of 3 × 10^5^ cells were plated in each well of a 6-well plate with 3 ml of DMEM and incubated for 24 h as described above. The cells were cotreated with GEM and/or phenothiazine for 3 days. After 3 days, the surviving cells were washed with 2 ml DPBS, detached with 200 μl of trypsin for 1-2 min, resuspended in fresh drug-free DMEM and incubated for 24 h. After 24 h, the cells were collected as described above, transferred to a microcentrifuge tube in PBS, and enumerated with trypan blue solution (0.4%) and the automated cell counter. If the surviving cell levels were under the limit of detection, we used a flow cytometer. To do this, the cell suspension was centrifuged at 800 rpm. for 5 min, and the supernatant was removed. The cell pellet was resuspended in 500 μl of 0.85% sodium chloride (NaCl) solution. Then, the cells were stained with 0.25 μM SYTOX green (catalog # S7020, Thermo Fisher Scientific) and SYTO60 red (catalog # S11342, Thermo Fisher Scientific) and then incubated at 37 °C for 15 min. SYTOX green is cell impermeant and only stains dead cells. SYTO60 is cell permeant and can diffuse through the cell membrane. After 15 min, the cell suspension was centrifuged at 800 rpm. for 5 min, and the supernatant was removed. Finally, the cell pellet was resuspended in 500 μl of 0.85% NaCl solution and transferred to a 5-ml test tube for flow cytometry analysis. The cells were excited at 488 nm for green fluorescence and 561 nm for red fluorescence. The green fluorescence was detected with a 520-nm emission filter; the red fluorescence was detected with a 615-nm emission filter.

### Statistical analysis

GraphPad Prism 8.3.0 was used for linear regression analysis, and the slopes of untreated and treated groups were compared with F statistics. The threshold of significance was set to P<0.05. Pairwise comparisons were performed using unequal variance t-tests or ANOVA with a significance threshold of P ≤ 0.05. A minimum of three independent biological replicates (unless otherwise stated) were assessed for all experiments. In all figures, data corresponding to each time point represent the mean value ± standard error.

## ACKNOWLEDGEMENT

We thank the members of the Orman Lab research group for their valuable contributions to this project. We also thank Barry Bochner and Brendan Lewis from Biolog, Inc., for providing us valuable insights and suggestions about our study. The research was supported by the University of Houston startup grant.

## AUTHOR CONTRIBUTION

P.K., V.A. and M. A. O. designed the study and interpreted the data. P.K. and V.A. performed the cell experiments. J.C. M. analyzed the transcriptomics data analysis. M. A. O. supervised the research and wrote manuscript with P.K., V.A., J. C. M. and M. A. O. contributed in reviewing the manuscript.

